# The Mechanical Influence of Densification on Initial Epithelial Architecture

**DOI:** 10.1101/2023.05.07.539758

**Authors:** Christian Cammarota, Nicole S. Dawney, Phillip M. Bellomio, Maren Jüng, Alexander G. Fletcher, Tara M. Finegan, Dan T. Bergstralh

## Abstract

Epithelial tissues are the most abundant tissue type in animals, lining body cavities and generating compartment barriers. The function of a monolayer epithelium – whether protective, secretory, absorptive, or filtrative –relies on regular tissue architecture with respect to the apical-basal axis. Using an unbiased 3D analysis pipeline developed in our lab, we previously showed that epithelial tissue architectures in culture can be divided into distinct developmental categories, and that these are intimately connected to cell density: at sparse densities, cultured epithelial cell layers have a squamous morphology (Immature); at intermediate densities, these layers develop lateral cell-cell borders and rounded cell apices (Intermediate); cells at the highest densities reach their full height and demonstrate flattened apices (Mature). These observations prompted us to ask whether epithelial architecture emerges from the mechanical constraints of densification, and to what extent a hallmark feature of epithelial cells, namely cell-cell adhesion, contributes. In other words, to what extent is the shape of cells in an epithelial layer a simple matter of sticky, deformable objects squeezing together? We addressed this problem using a combination of computational modeling and experimental manipulations. Our results show that the first morphological transition, from Immature to Intermediate, can be explained simply by cell crowding. Additionally, we identify a new division (and thus transition) within the Intermediate category, and find that this second morphology relies on cell-cell adhesion.

## Introduction

Epithelial tissues are cohesive cell sheets that serve to mechanically and compositionally protect animal body compartments. Epithelia most commonly form a monolayered architecture, with component cells adhering closely to their lateral neighbors and sharing a common apical-basal polarity axis (directionality) (Buckley and St Johnston 2022). Epithelial tissue function relies on tissue architecture, and a disruption of this architecture can lead to disease. For example, the most common form of adult cancer is solid carcinoma, which results from epithelial overproliferation and subsequent architectural disruption (McCaffrey and Macara 2011).

A full understanding of how cells form the shapes required for layered epithelial architectures remains elusive. While the molecular effectors of apical-basal polarity have been widely studied, the physical basis for cell shape in the apical-basal axis of tissues is relatively unexplored. In a recent study we used an unbiased image analysis pipeline called ALAn (Automated Layer Analysis) to measure apical-basal architecture in Madin-Darby Canine Kidney (MDCK) cultured epithelial cell layers (Dawney et al. 2023). ALAn recognizes a developmental series of three organized architectures (Immature, Intermediate, and Mature), and a fourth category of exclusion (Disorganized) (Dawney et al. 2023). The organized apical-basal architecture types we identified are distinguished as follows: 1) Immature layers are composed of flat cells which do not form significant cell-cell contacts; 2) Intermediate layers are taller and have well defined cell-cell contacts; 3) Mature layers are the tallest organized layers, well defined cell-cell contacts and have a flattened apical surface (Figure 1, Figure S1). Architectures transition through the developmental series as density increases, and each architecture is associated with a particular density regime (Figure 1).

**Figure 1:**
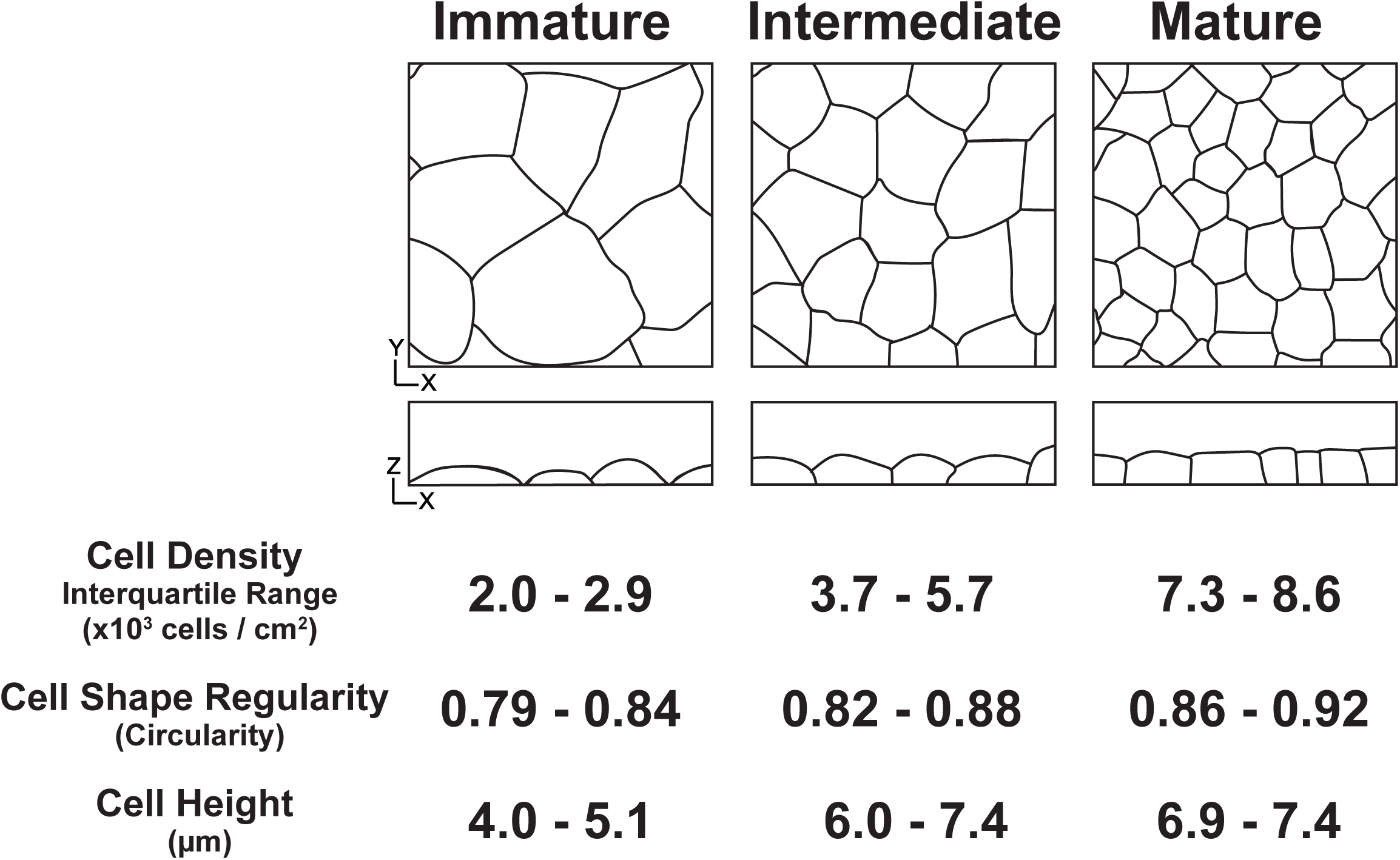
MDCK cells form a developmental series of architectures in culture. Cartoons summarizing the distinct epithelial architecture categories made by cultured epithelial cells along with their defining average density, regularity and height values (defined in Dawney et al., 2023).

These three organized apical-basal architectures correspond to changes in cell shape and arrangement with respect to the apical tissue surface. Across metazoans, epithelial cells in a proliferating tissue are chiefly 5-, 6-, and 7-sided polygons at the apical surface, with the plurality being hexagons (Thompson 1917; Farhadifar et al. 2007; Gibson et al. 2006). One explanation for this observation is that hexagonal packing represents a minimum energy state (Thompson 1917; Honda and Eguchi 1980). Because cell shapes and cell packing are intertwined, cell shape regularity (as measured by circularity) can be used to reflect packing efficiency. Apical cell shapes, and therefore packing, become more regular as cultured cell layers transition through the developmental series (Figure 1, Figure S1).

The Disorganized architectural category we identified in culture is characterized by a failure to achieve a regular monolayered architecture in the apical-basal axis, and occurs when the underlying cell culture substrate area is insufficient to accommodate the number of cells seeded onto the plate (Dawney et al. 2023). On the basis of this observation we previously proposed a simple conceptual model; layer organization is determined by a competition between cell-cell and cell-substrate adhesion (Dawney et al. 2023). In the absence of available substrate, cell-cell adhesion wins the competition and monolayering is not achieved. While monolayer formation is a mechanical consequence of adhesion in this simple model, subsequent steps in architecture development are not accounted for.

In this study we used a combination of computational modeling and cultured-cell experimentation to ask whether the developmental transitions from one organized architecture to the next are mechanically driven. Given that the mechanical parameter most obviously associated with these transitions is density, we focused in particular on the question of whether these transitions are driven by crowding. We find that crowding can explain the first transition (Immature to Intermediate) but not the second, and that the initial appearance of Intermediate architecture is not restricted to epithelial cells. However, we also identify a new transition within the Intermediate category. Our findings indicate that this second transition is the onset of a strictly epithelial architecture.

## Results

### Development of a 2D Multinodal Cell Model

Computational models, in particular vertex models, have been used to simulate epithelial cell shapes with respect to the tissue plane (Fletcher et al. 2014). We took a similar approach to address how mechanical forces impact cell (and therefore tissue) shape in the apical-basal plane. We developed a discrete biophysical model of cell shape dynamics in the apical-basal plane, similar to previous ‘deformable polygon’ models (Tamulonis et al. 2011; Perrone, Veldhuis, and Brodland 2016; Boromand et al. 2018).

Cell cortices and the substrate are each discretized into a set of nodes or ‘interaction sites’ that line the cortex of an XZ cellular cross section (Figure 2). Each cell is subject to the following forces: external adhesions (cell-cell and cell-substrate); internally generated forces regulating size and shape via cytoskeletal dynamics; and an active spreading force driven by actin-based lamellipodia (Figure 2). In addition to these forces, a feedback mechanism allows cell-cell and cell-substrate adhesions to strengthen each other. Cell nodes are subject to a rigid substrate; this is implemented by preventing any node from going below the z = 0 line.

**Figure 2:**
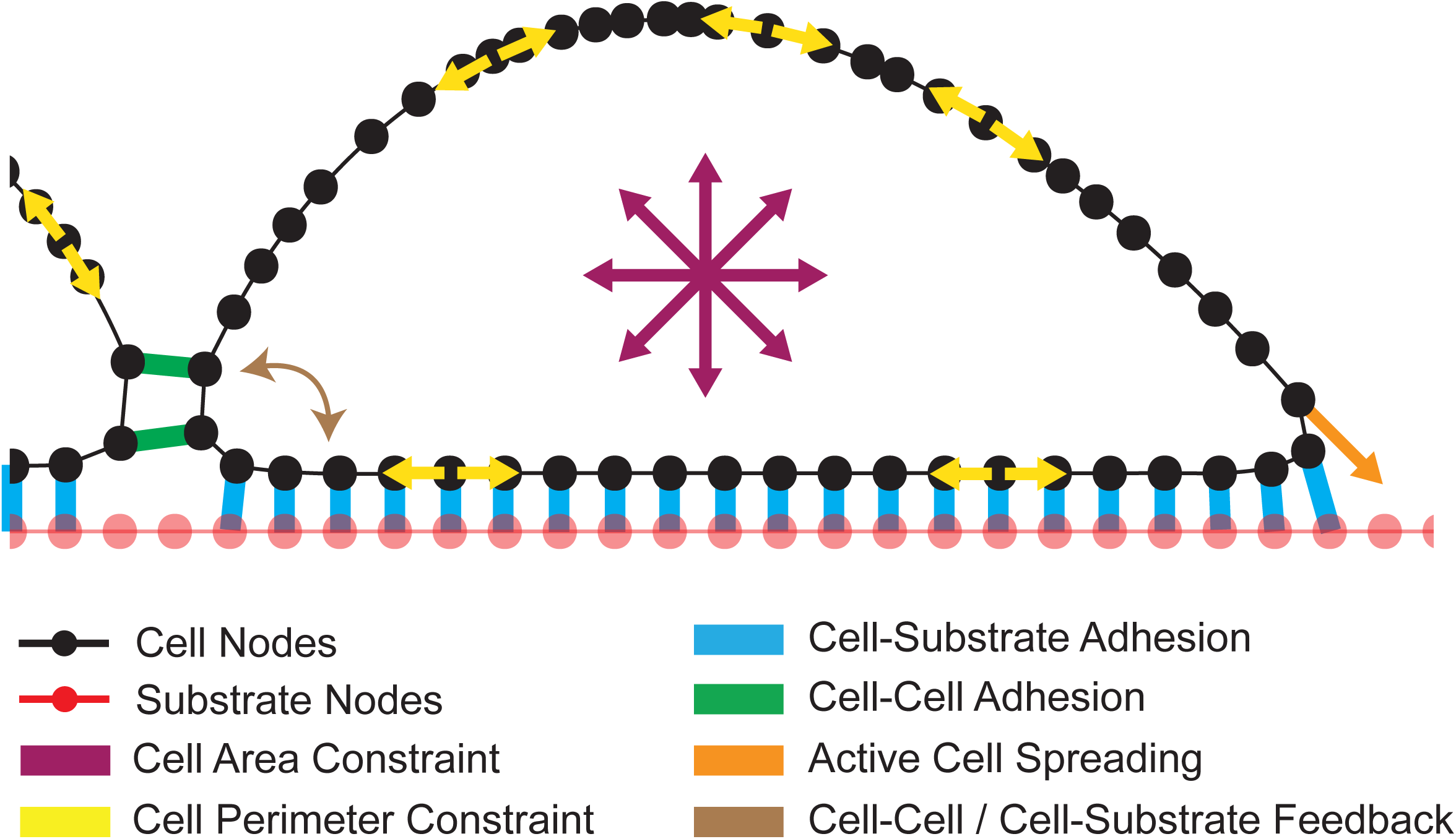
Introduction to the computational cell model. Schematic representing the forces accounted for in the computational model.

Cells remodel their cortex; if the spacing between adjacent nodes surpasses 2 times the expected length, a new node will be added at the bisection. If the spacing between adjacent nodes becomes smaller than half the expected length, the node which is closest to its next nearest neighbor will be removed. Remodeling in this way keeps the cells at a roughly constant interaction point density throughout a simulation.

External adhesion is modeled by springs that link interaction sites when they come into proximity. The spring breaks when connected nodes move outside of a threshold interaction distance. Cell-substrate adhesions are calculated using:

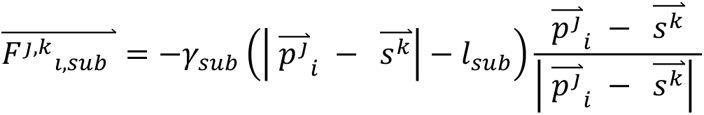

where *F^j,k^_i,sub_* is the force on the j^th^ node of the i^th^ cell due to adhesion to the k^th^ substrate point which points from 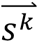 to 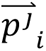, *γ_sub_* controls the adhesion strength between cells and the substrate, 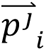 is the position of the j^th^ node of the i^th^ cell, 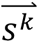 is the position of the k^th^ substrate point, and *l_sub_* is the natural length of a cell substrate adhesion. This force only exists when) 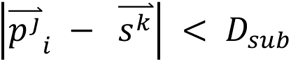 where *D_sub_* is the maximum interaction distance for a substrate adhesion. Substrate interaction sites will make a connection to only the closest interaction site on a cell, and each cell interaction site can only make one substrate connection. Similarly, cell-cell adhesions are calculated using:

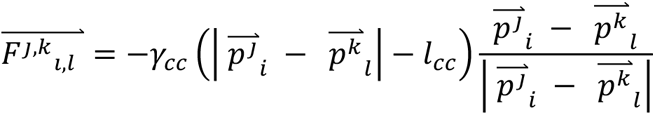

where *F^j,k^_i,l_* is the force on the j^th^ node of the i^th^ cell due to adhesion to the k^th^ node on the l^th^ cell (which points from 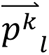 to 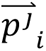), *γ_cc_* controls the adhesion strength between cells, 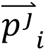 is the position of the j^th^ node of the i^th^ cell, 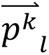 is the position of the k^th^ node of the l^th^ cell, and *l_cc_* is the natural length of a cell-cell adhesion. This force only exists when 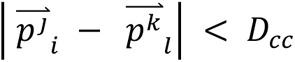 where *D_cc_* is the maximum interaction distance for a cell-cell adhesion. When 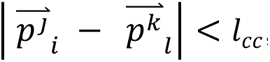, we replace *γ_cc_* with a repulsive strength of 10 N/m.

The feedback mechanism between cell-cell and cell-substrate connections operate as follows: if a cell is connected to the substrate, *γ_cc_* is increased by 3% for each substrate connection; if a cell is connected to at least two other cells, *γ_sub_* is increased by 50%; if a cell is connected to at least one other cell, *D_cc_* is increased by 20%.

Internal regulation of cell size and shape is modeled by having each cell maintain a constant area in XZ while minimizing the cortical perimeter. The internal forces are governed by the following energy equation:

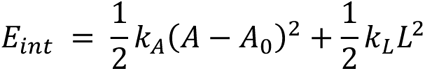

where *k_A_* controls the constraint on the cross-sectional area of the cell, A is the current cross-sectional area, *A*_0_ is the constant, natural cross-sectional area, *k_L_* controls the constraint on the perimeter of the cell and L is the current perimeter of the cell. We take the negative gradient of the above energy with respect to nodal coordinates to calculate the internal forces on each node. Cells tend to round up because of internal regulatory forces in both the model and in culture.

Gravity is estimated to be four to six orders of magnitude smaller than the other forces in this system, and is therefore excluded from consideration once cells reach the substrate or a cell connected to the substrate, as expected in a low Reynolds’ number environment (Purcell 1977). The parameter values in my simulation are set by the estimate of ∼100 µm^2^ XZ cross section for MDCK cells, and reported values of physical forces for cultured cells (Brückner and Janshoff 2015; Gallant, Michael, and García 2005; Chu et al. 2004).

### Implementation of Active Spreading in the Model

We first attempted to model the behavior of a single cell in contact with a substrate using the given implementations. Our initial expectation was that cell-substrate adhesion would be sufficient to promote the flat shape of isolated cells in culture (Figure S2A), but we found this not to be the case; simulated cells maintain a round shape under the influence of substrate adhesion alone (Figure S2B). In culture, spreading relies on active remodeling (protrusion) of the cytoskeleton in addition to adhesion (Pierres, Benoliel, and Bongrand 2002; Chamaraux et al. 2005; Cuvelier et al. 2007). We therefore implemented an active cell-spreading component to our simulations, introducing a constant force (1.6e-5 N), pointing 45° outward and below horizontal that acts on nodes immediately adjacent to the outermost substrate connections. Active spreading initiates once 10% of all nodes contact the substrate. The addition of an active spreading force recapitulated the cell shapes observed in culture (Figure S2C).

We also incorporated contact inhibition of locomotion, a phenomenon observed across various systems, into our model (Roycroft and Mayor 2016). To achieve this, the spreading force drops to zero when a simulated cell contacts a neighbor. Contact inhibition is implemented at each edge of the cell independently, meaning that spreading continues on the other side of the cell until/unless it also contacts a neighbor.

The refined model recapitulates our live cell imaging. Both cultured and simulated cells tend to maximize substrate connections at the expense of cell-cell adhesions, and a simulation of two cells in proximity closely reflects the same configuration of cultured MDCK cells (Figure 3A).

**Figure 3:**
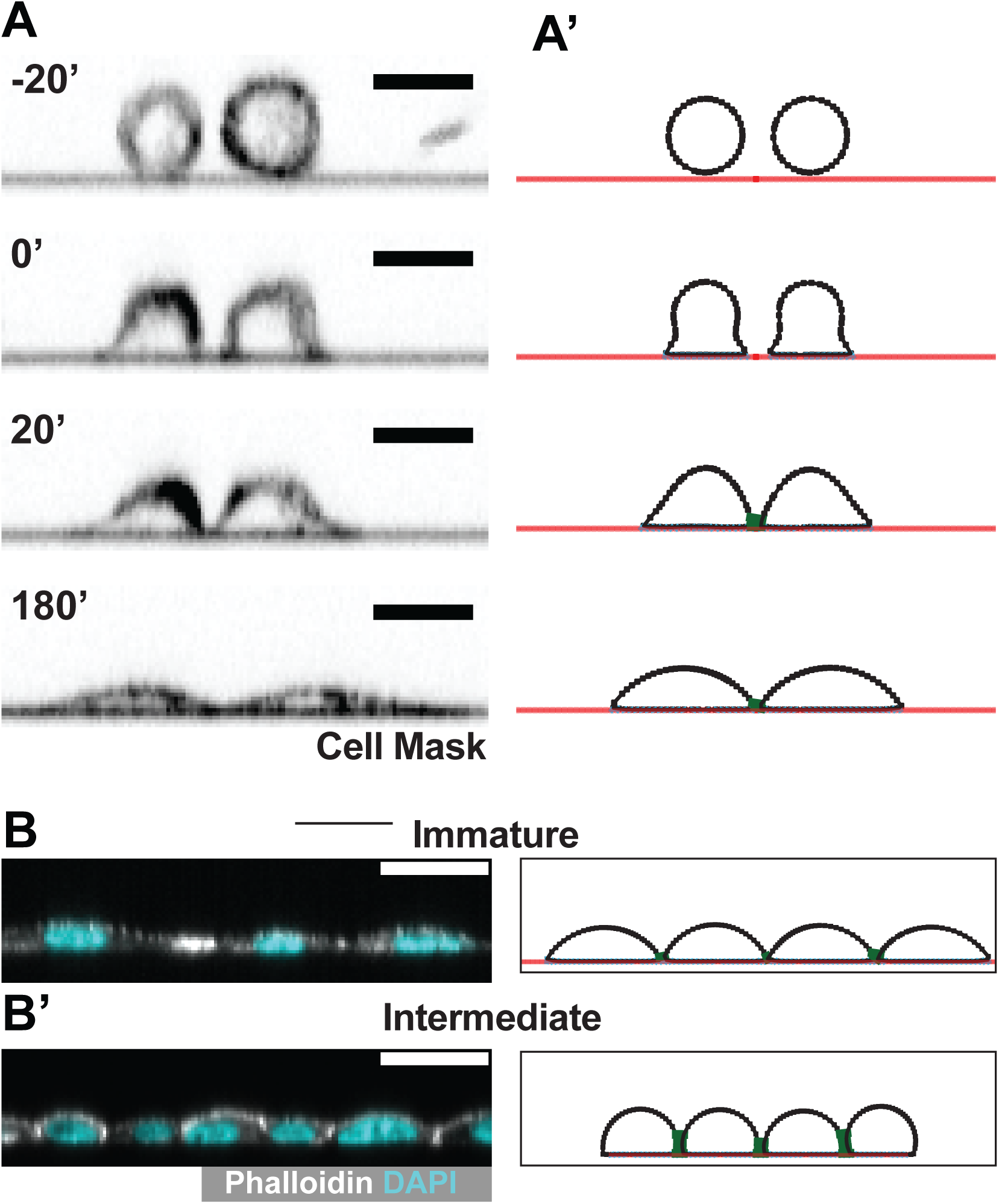
The computational model recapitulates the behavior of MDCK cells in culture. Side by side comparison of live MDCK cells (left) and simulated ‘cells’ (right) show that the model can recapitulate real cell behavior in the contexts of: A) cell ‘plating’-dropping cells onto a substrate, and B) layer densification, and therefore epithelialization. Simulated cells exhibit Immature architecture at low densities, and Intermediate architectures at higher densities. Scale bars = 20 µm.

### Intermediate Epithelial Architecture Arises as Simulated Cells Densify

Cultured monolayer architecture changes as the layer densifies (Figure 1) (Dawney et al. 2023), suggesting the possibility that epithelial architecture arises from cell crowding. We used our computational model to test this possibility, varying cell density by limiting the amount of available basal substrate. As a 2D model of a 3D system, density cannot be directly compared between the simulated and physical layers. We therefore report simulated cell density in terms of arbitrary units (Arb.), derived by scaling the linear cell density. At essentially unlimited substrate, cells spread without developing many contacts and cells form architectures resembling Immature architecture (Figure 3B). Lateral surfaces develop as the available substrate is decreased (Figure 3B’). These surfaces are observed in our simulations when 10% of all nodes are dedicated to cell-cell adhesion, and we therefore use this point to define the transition from Immature Intermediate architecture.

The flat apical surface characteristic of Mature layers is not recapitulated in our simulations even at the highest densities. We confirmed this by comparing the ratio of apical length to basal length in MDCK cells and simulated cells. In both cases, this ratio increases gradually over the densities associated with Immature and Intermediate architectures. In the simulated cells, a gradual decline begins at a density near 7 Arb (Figure S3). In MDCK cells the ratio demonstrates a steeper decline beginning at ∼7×10^3^ cells/mm^2^, correlating with the appearance of flattened apices (Figure S3).

Our simulations recapitulate the development of Immature and Intermediate, but not Mature, architectures. These results suggest that the appearance of Intermediate architecture is a straightforward mechanical consequence of deformable objects crowding together. Contrastingly, the transition from Intermediate to Mature architecture relies on an additional biophysical step(s) initiated by densification.

### Division Pressure Drives the Immature to Intermediate Transition

Our modeling and culture data predict that cell density is the driving force behind the emergence of Intermediate epithelial architecture. This raises the question of what drives densification in epithelia. The most obvious answer is cell division, and we therefore implemented division in our computational model.

We simulated growth of a three-cell colony along an unbound substrate, “seeding” it such that a central cell divides before its neighbors. After this cell divides, the two daughter cells develop multiple cell-cell contacts with each other and their neighbors, reflecting Intermediate architecture (Figure 4A). At this point the colony edge is pushed outward, in agreement with earlier work showing that cell colony growth follows the Fisher-Kolmogorov equation (Marel et al. 2014; El-Hachem et al. 2019), which predicts that a proliferative colony will exhibit a cell density gradient with its highest point at the center.

**Figure 4:**
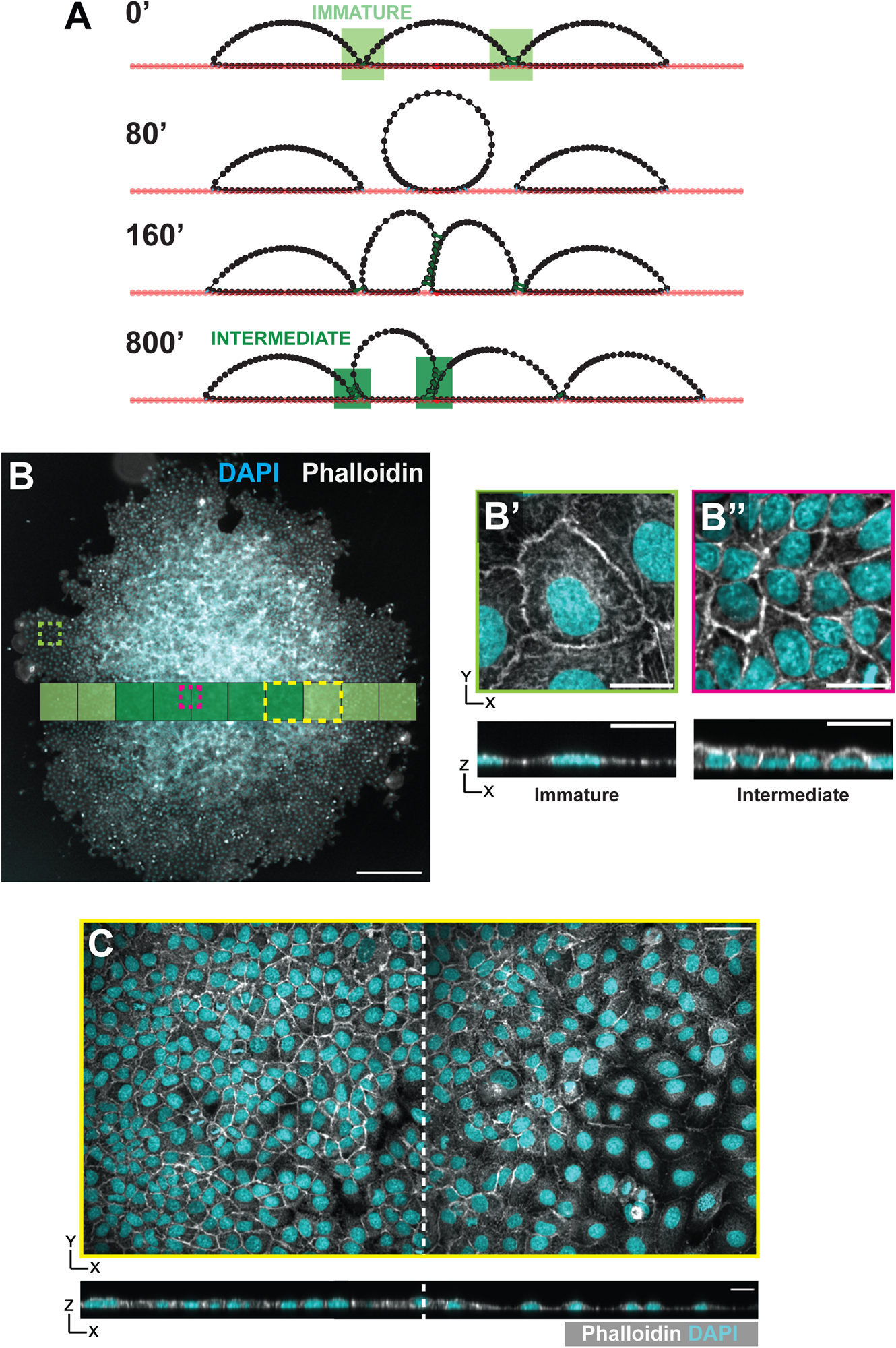
Division pressure drives the Immature to Intermediate architectural transition. A) Timepoints from a simulation of a 3-cell layer with a division implemented in the central cell. Division leads to an increase in cell-cell density at the center of the monolayer, and a gradient of cell-cell contacts from the edge to the center of the layer. B) An isolated cell colony of MDCK cells grown from sparse cell seeding onto the culture substrate. The colony exhibits a density gradient from the center to the edges as predicted by the model. Cells at the edge of the colony form an Immature architecture (B, and B’) whereas cells in the dense central region form an Intermediate architecture (B and B”). C) High resolution imaging of the region highlighted in the yellow box from B shows the gradient in cell density. Scale bar in B = 500 µm. All other scale bars = 20 µm.

Both theory and our simulations predict that division pressure, the force generated by cell division in a tissue, promotes the transition from Immature to Intermediate architecture. We tested this prediction by culturing MDCK cells at a low density, creating colonies of cells that were smaller than the growth area of our culture plates and therefore unconfined at their edges (Figure 4B). Our MDCK colonies clearly showed a density gradient from the center to the edge of the colonies as predicted (Figure S4). Using our image analysis pipeline (ALAn) (Dawney et al. 2023), we found that layer architectures are Immature at the colony edge and Intermediate at the colony center (Figure 4B,C). These results agree with previous studies showing that cells in the central region of a growing colony demonstrate greater shape regularity (with respect to the tissue surface) than cells at the edge, as expected from the transition from Immature to Intermediate architecture (Puliafito et al. 2012; Metzner et al. 2018; Khain and Straetmans 2021). Together these findings suggest that densification is sufficient to drive the transition from Immature to Intermediate architecture.

### Intermediate Architectures Develop When Cell-Substrate Adhesion and Spreading are Impaired

Having determined that Intermediate architecture development relies fundamentally on crowding, we next set out to investigate the role played by cell-substrate adhesion. *In vivo,* epithelia are anchored in position by interaction with a basement membrane rich in extracellular matrix (ECM) (reviewed by (Kozyrina, Piskova, and Di Russo 2020).). The basal substrate provides physical and biological networks that regulate cell behavior and tissue architecture (Maruthamuthu et al. 2011). This is modelled in culture by providing ECM on the cell culture substrate and is necessary for epithelialization (Wang, Ojakian, and Nelson 1990).

To model the development of Intermediate architecture *in silico,* we time-evolved a simulated four-cell colony for a period corresponding to 24 hours in culture. We restricted our modeling to densities below 7 Arb (Figure S3). We performed simulations over a range of cell-substrate adhesion values, starting with 20 N/m, which is physiologically relevant and used as our standard (Gallant, Michael, and García 2005). The relationship between cell-substrate adhesion (as mediated by matrix receptors) and active protrusion is difficult to parse experimentally, but we and others observe that limited MDCK cell spreading occurs even on uncoated glass (Leighton et al. 1970) (Figure S5A). We therefore undertook simulations using three models for active spreading: 1) the spreading force is held constant at all substrate adhesions; 2) the spreading force scales non-linearly with substrate adhesion; 3) the spreading force scales linearly with substrate adhesion. We consider the first condition to represent an upper bound and the third to represent the lower bound, whereas the second most closely represents our observations in culture.

We used the proportion of total interaction points (per cell) that participate in cell-substrate or cell-cell adhesion as proxies for cell-substrate interface and cell-cell interface length. These parameters normally decrease and increase (respectively) as an MDCK layer matures (Figure S5B,C). As expected, we found that reducing cell-substrate adhesion strength reduces the number of cell-substrate connections (Figure 5A, Figure S5D,F). Somewhat counterintuitively, it also reduces the number of cell-cell connections, especially at higher densities (Figure 5A’, Figure S5D’,F’). This is because spreading in our model is outcompeted by the internal regulation of cell shape, which tends towards circularity at weaker cell-substrate adhesion strengths (Figure 5B, Figure S5E,G).

**Figure 5:**
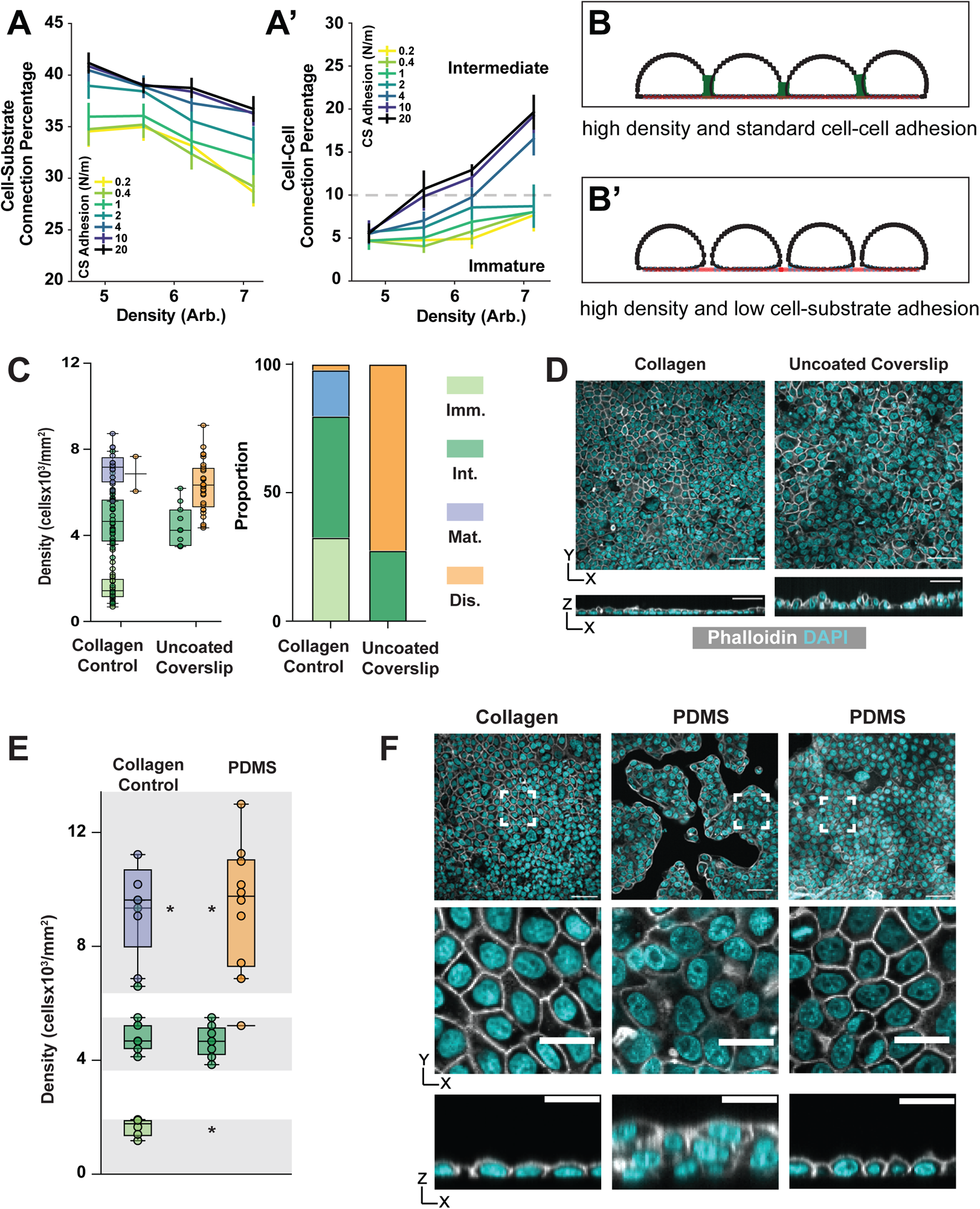
Intermediate architecture can form despite decreased cell-substrate adhesion. A) Cell-substrate connections decrease (A) and cell-cell connections increase (A’) as cell density increases across tested values of cell-substrate adhesion strengths when spreading is scaled nonlinearly in simulations. The dashed line represents the connection percentage required for Intermediate architectures to arise in the model (A’). B) Model representative displaying the final result from decreasing cell-substrate adhesion strengths. C and D) Plating cells on uncoated coverslips leads to primarily Disorganized architectures, but Intermediate architectures can still be found. Representative images shown in D. Scale bar = 50 µm. E and F) Plating cells on a PDMS membrane leads to islands of primarily Disorganized architecture but Intermediate architectures are still found (E). Representative images shown in F. Highlighted areas in E represent density cut-offs at which different architectures should be observed. Due to the presence of islands, smaller areas of each image are analyzed (dashed boxes, F) at the densities of which each layer type should be observed.

Spreading strength modulates the role of substrate adhesion in the transition from Immature to Intermediate architecture in our simulations. In both the nonlinear and upper bound spreading conditions, Intermediate architectures develop even when cell-substrate adhesion is reduced by as much as 80% (from 20 N/m to 4 N/m) (Figure 5A’, Figure S5D’). In the lower bound condition, in which spreading and adhesion have a linear relationship, Intermediate architectures are observed at 50% adhesion strength (Figure S5F’). Thus, while Intermediate architectures develop in each spreading mode, the amount of substrate adhesion required varies inversely with the spreading strength.

Our simulations show that Intermediate architectures can develop when spreading and substrate adhesion are reduced. We tested this in culture by changing the substrate. The standard culture wells used in this and our earlier study are coated with collagen, an ECM component that provides a basis for receptor-mediated adhesion but is not required for MDCK cell adherence to the culture well, meaning that adhesion is diminished but not lost in its absence (Leighton et al. 1970). We shifted to uncoated wells and found that the layer architecture profile changed dramatically:

Disorganized layers predominated at the densities we expect to find Mature layers (> 5×10^3^ cells/mm^2^), and no Immature architectures were observed. (Figure 5C,D). Both observations are predicted by our conceptual model for monolayer development, which is that monolayers rely on a competition between cell-cell and cell-substrate adhesion. In the collagen-free condition, the balance is expected to shift towards cell-cell adhesion, meaning that cells will tend to accumulate (clump) rather than cover the substrate. At high densities this shift causes cells to pile up, leading to Disorganization. At low densities it prevents confluence. In agreement with the latter, approximately 20% of regions imaged on the uncoated substrate were subconfluent.

Roughly 30% of the layers analyzed demonstrated the characteristics of Intermediate architecture, both with respect to apical-basal and apical (surface) cell shapes. These layers also fell in the expected ranges for cell density (Figure 5C) and cell shape regularity with respect to the surface (Supplemental Figure 5H). This result agrees with our simulations; Intermediate architectures can develop despite reduced cell-substrate adhesion.

We also plated cells on a biologically inert polydimethylsiloxane (PDMS) membrane. This substrate not only lacks collagen, as in the previous experiment, but is also less stiff (∼1 MPa for our membrane from ∼1GPa for our coverslip) and should therefore curtail active spreading (Larson, n.d.; Discher, Janmey, and Wang 2005; Yeung et al. 2005). In agreement, cells plated on PDMS do not spread to cover the available substrate, as they do on collagen, but instead form “islands” with free space in between. This presented an obstacle for analysis. Our standard images are ∼300 µm in the X and Y dimensions, but only 3 of the 35 regions analyzed in this experiment were confluent over that area. To overcome this obstacle we reduced the area of analysis to 60 µm x 60 µm (Figure E,F). This introduced a bias to the experiment, since areas of analysis had to be determined by the experimenter. To limit this bias we restricted the analysis to those density regimes most closely associated with the three organized architectures (Figure 1). As observed on the uncoated coverslips, Disorganized architectures predominate at densities above 5×10^3^ cells/mm^2^ and Immature architectures do not develop. Intermediate architectures can be observed at their expected density regime.

Taken together, our modeling and experimental results support a mechanical explanation for the transition from Immature to Intermediate architecture. They also suggest that densification is the main driver of this transition, whereas cell-substrate adhesion and active cell spreading, a parameter not accounted for in our earlier conceptual model for monolayering, play a less important role.

### Cell-Cell Adhesion Facilitates the Transition From Immature to Intermediate Architecture but is not a Major Factor

MDCK cells A) express the cell-cell adhesion factor E-cadherin, which has long been associated with epithelial identity, and B) are one of the few cultured cell types that effectively epithelialize (Behrens et al. 1985; Nelson et al. 1990). We therefore investigated the question of how cell-cell adhesion contributes to the transition from Immature to Intermediate architecture. Starting with our standard cell-cell adhesion strength of 0.2 N/m – a value determined empirically for cultured cells expressing E-Cadherin (Yeh-Shiu Chu et al 2004) - we modulated the strength of cell-cell adhesions in our simulations by two orders of magnitude in either direction and determined cell-substrate border length and cell-cell border length. We found that while reducing cell-cell adhesion strength does not prevent the development of Intermediate architecture in our simulations, the density at which Immature architecture transitions to Intermediate (dashed line in Figure 6A’) increases (3.1×10^3^ cells/mm^2^ in our control versus 3.6×10^3^ cells/mm^2^ when adhesion is reduced to 0.002 N/m) (Figure 6A). These results suggest that cell-cell adhesion facilitates the development of Intermediate cell shapes at lower densities (<∼4×10^3^cells/mm^2^), but is not an absolute requirement.

**Figure 6:**
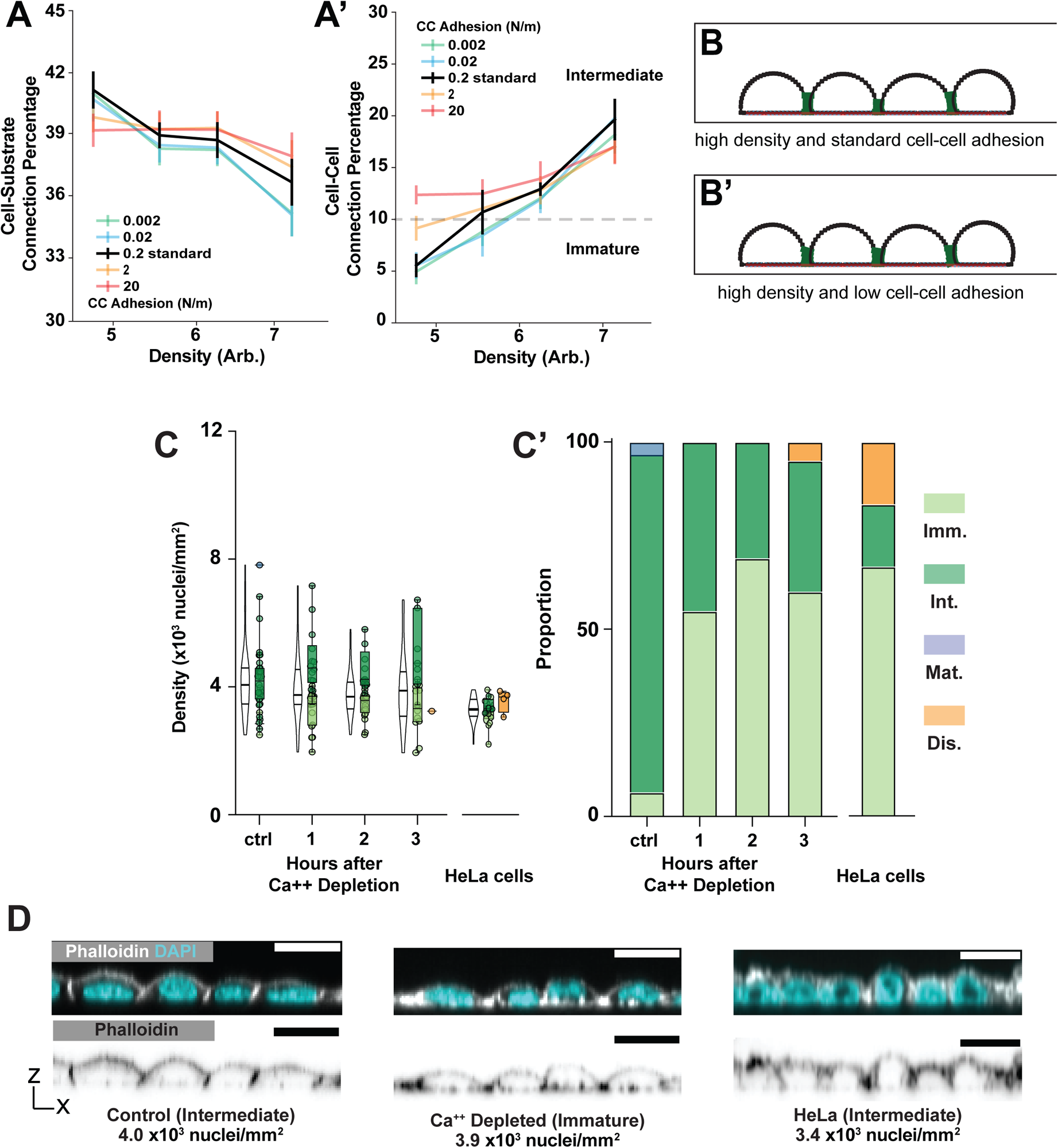
Intermediate monolayers form with decreased cell-cell adhesion. A) Cell-substrate connections decrease (A) and cell-cell connections increase (A’) as cell density increases across tested values of cell-cell adhesion strengths in simulations. A’) The dashed line represents the cell-cell connection percentage required for Intermediate architecture to arise in the model. B) Model representative displaying the final result from decreasing cell-cell adhesion strengths. C, C’) Calcium depletion leads to Immature layer architectures in the low density regime where Intermediate architectures are expected. HeLa cells develop Immature, Intermediate and Mature architectures. D) Representative images show differences in morphology between control and calcium depleted layers at comparable densities. Representative HeLa cells shown with an intermediate morphology.

To test this prediction, we allowed MDCK cells to develop Intermediate architectures under standard conditions (200K cells plated for 24Hr in a 1 cm^2^ culture area), then replaced the culture medium with a calcium-free medium to disrupt calcium-dependent cell adhesion (cadherins) (Takeichi 1977). This resulted in a retrograde shift in layer architecture - from Intermediate “back” to Immature - but only at the lowest densities at which Intermediate architectures develop (≤ 4×10^3^ nuclei/mm^2^) (Figure 1, Figure 6C). A potential caveat to interpretation is that calcium depletion might also impact cell shape by reducing non-muscle myosin II activity, and therefore cell contractility (Szent-Györgyi 1975) We used Blebbistatin, which inhibits non-muscle myosin II directly, to test this. Blebbistatin had no significant impact on architecture, indicating that the effect of calcium depletion is most likely a consequence of reduced cell-cell adhesion (Figure S6A).

As a second approach we examined architecture development in HeLa (Henrietta Lacks) cells, which do not express E-cadherin (Figure S6B) (Vessey et al. 1995). We cultured these cells to 3.6-4×10^3^ nuclei/mm^2^ (their maximum density in our laboratory) using the same collagen-coated culture wells used for our MDCK cell experiments. We suspect that this density does not correspond directly to those observed for MDCK cells because HeLa cells are larger, in which case they should be expected to achieve comparable crowding at a lower cell density (X. Liu, Oh, and Kirschner 2022). In addition to the expected Immature architectures, we detected Disorganized and, at higher densities, Intermediate architectures (Figure 6C). Together our modelling and experimental work show that adhesion facilitates the transition from Immature to Intermediate architecture at low densities but is not necessary for this transition at higher densities.

### Some Intermediates are More Equal Than Others

Our results show that the appearance of lateral cell surfaces – our definition for the transition between Immature and Intermediate – is not restricted to epithelial cells in culture. This led us to ask the question of how well the Intermediate category recognized by ALAn reflects epithelial architecture. To address this question, we examined the development of another epithelial characteristic, namely cell shape regularity with respect to the tissue surface, as cells densify. Across our large control data set we find that the relationship between density and circularity (shape regularity) is linear up to a circularity of ∼0.84, at which point it begins to plateau (Figure 7A).

**Figure 7:**
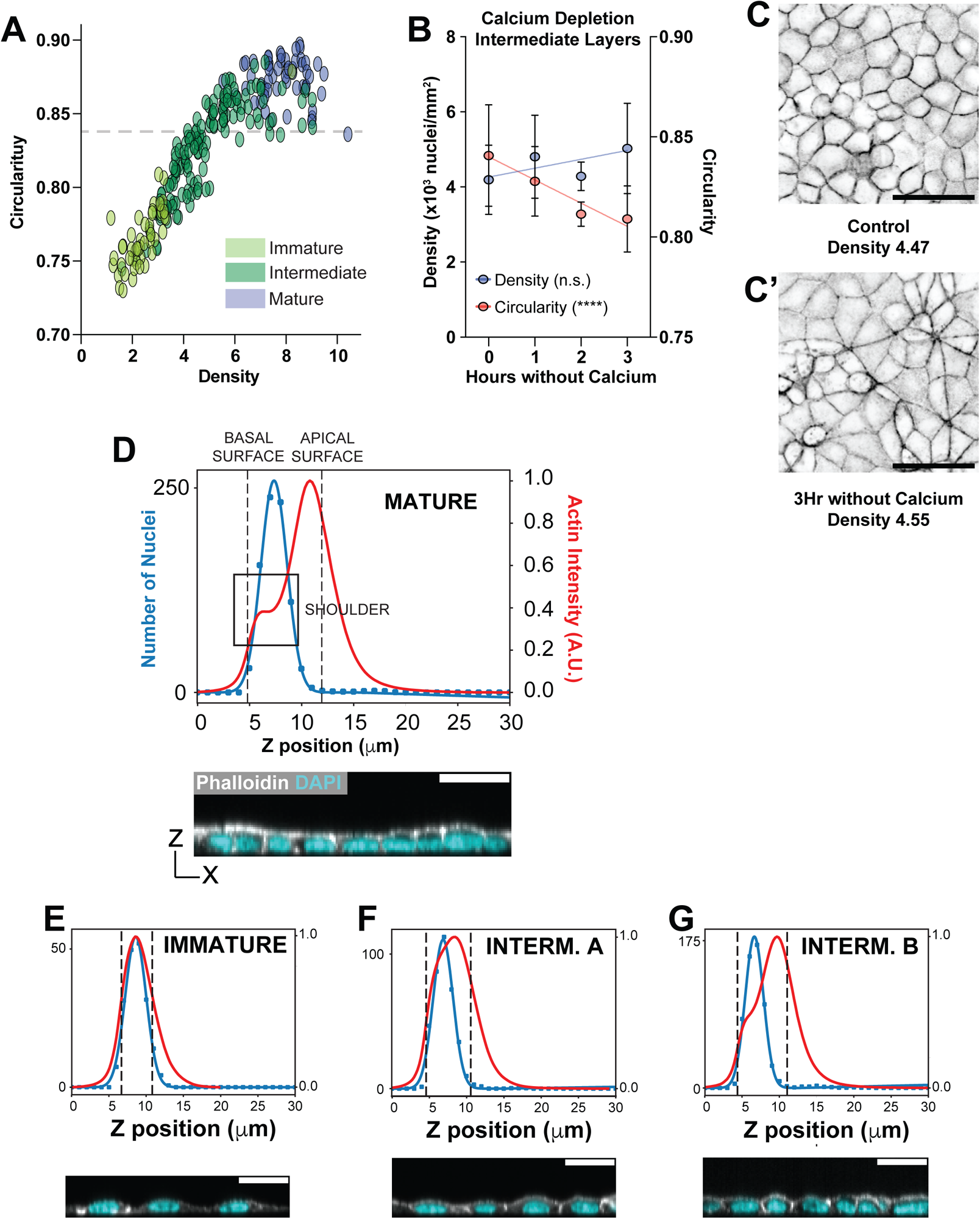
Intermediate epithelial architecture can be divided into two sub-categories. A) Density and cell shape regularity (circularity) have a linear relationship until a circularity of approximately ∼0.84, at point circularity plateaus. B). Cell shape regularity decreases in MDCK cells as a function of hours without Calcium. The slope has a p < 0.0001 to be non-zero. Significance was determined using an F-Test. Density of analyzed layers is unchanged as a function of hours without Calcium. The slope has a p = 0.0651 to be non-zero. Significance was determined using an F-Test. C) Representative images showing the change in cell shape (with respect to the tissue surface) after calcium depletion. D) ALAn-generated Nuclei and Actin distribution plots across the apical-basal depth of a tissue layer, formed by projecting Actin onto the z axis, and making a histogram of segmented nuclei positions. This type of plot was developed to categorize epithelial architectures in Dawney et al (2023). A distinct ‘shoulder’ is present in the plot of actin density against z position in Mature epithelial layer types, as indicated by the box, corresponding to distinct pools of lateral and apical actin. ALAn uses the derivative of the actin intensity plot to detect this shoulder. If the derivative is two-peaked, the ratio of the right peak to the left peak is used. A ratio of greater than 1 is classed as a Mature layer. E,F and G) We now introduce a further sub-categorization of Intermediate layers based on the projected actin and nuclei profiles of layers across the apical-basal axis. Immature layers are characterized by a lack of defined lateral surfaces. Therefore, in these layers the peak nuclear distribution is located at, or occasionally above, the peak actin intensity (E). Intermediate layers develop a lateral surface and an asymmetric Actin intensity that peaks closer to the surface. Intermediate ‘A’ layers do not develop a clear shoulder (F). Intermediate ‘B’ layers develop a clear shoulder with a Lateral to Apical Shape Index of less than 1 (G).

We find that circularity decreases after calcium depletion, from an average of ∼0.84 in the control to ∼0.81 at three hours without calcium, and that this is not due to a change in cell density (Figure 7B,C’). Consistent with this, HeLa cells at high densities do not achieve the polygonal cell shapes associated with epithelia, but rather retain their characteristic spindle-shaped morphology even when their apical-basal architecture is recognized as Intermediate (Figure S7A). These findings show that shape regularity is not simply proportional to cell density. Instead, it is proportional to a sum of density and adhesion, taking the form: Circularity ∝ (α ∗ Density + β ∗ Adhesion), where α and β represent the relative contribution of Density and Adhesion to circularity respectively.

Together, our observations suggest that a uniquely “epithelial” architecture (relying on cadherin-mediated adhesion) corresponds to a shape regularities of ∼0.84 and above. We next asked whether layers below and above the 0.84 breakpoint could be distinguished in respect to their apical-basal architecture. Our analysis pipeline, ALAn, discerns the shape of lateral and apical surfaces at multi-cell scale using a profile of actin intensity projected along the apical-basal tissue axis (examples Figure 7D-G). A “shoulder,” corresponding in position to lateral actin at cell-cell borders, is evident in the actin intensity profile of a Mature layer (Figure 7D). The basal side (left in the profile) of the shoulder becomes evident as lateral surfaces develop, and the apical side (right in the profile) as apical cell surfaces flatten. The ratio between the maximum slopes on either side of the shoulder, defined here as the Lateral-to-Apical Shape Index, is therefore an indicator of architecture development. Our image analysis pipeline ALAn defines Mature architectures as those in which the Lateral-to-Apical Shape Index is greater than or equal to 1 (Dawney et al. 2023).

ALAN does not use the Lateral-to-Apical Shape Index to distinguish Intermediate architectures from Immature architectures. Rather, these are distinguished by A) an apically-biased asymmetry in the actin intensity plot and B) a change in the relationship between peak actin intensity and peak nuclear distribution across the apical-basal axis, such that the former is apical to the latter (meaning that an apical surface has started to develop in the cultured tissue) (Dawney et al. 2023). We examined how this Index evolves in early architectures, and whether it could be used as a means of assessing architecture development within the Intermediate group. We find that in the majority of Intermediate layers (80/141) in a large control data set, the Index cannot be determined. This is because the basal side of the shoulder develops before the apical side, indicating that lateral surface development precedes apical surface flattening. Nearly all (74/80) of these layers have an apical surface circularity below 0.84. Of remaining 61 Intermediates, in which ratio can be determined, 52 are above 0.84. On the basis of these findings, we divide Intermediate apical-basal architectures into two categories, Intermediate A (IntA, at an interquartile density range of 3.4-4.52 x 10^3^ cells/cm^2^) and Intermediate B (IntB, at 5.2-6.92 x 10^3^ cells/cm^2^). IntB layers have Lateral-to-Apical Shape Indices and cell shape regularities associated with cadherin-mediated cell-cell adhesion, whereas IntA layers do not. Thus we have defined a novel morphological transition in cultured epithelia. Our computational model, which implements simple biophysical parameters, does not explain this transition.

## Discussion

### Limitations of the Study

Our experimental manipulations in culture suffer from a lack of precision - we have effectively used blunt tools (e.g. calcium depletion) on incompletely-identified fasteners:

- Both cell lines used in our study are notoriously heterogenous between labs (Dukes, Whitley, and Chalmers 2011; Y. Liu et al. 2019). We derived the “standard” adhesion strengths in our computational model from experimental results in the literature, but cannot be confident that these are accurate to our cell type.
- Calcium depletion will abrogate cadherin-based adhesions. We do not know the full adhesion profile of the MDCK cells in our lab, or to what extent non-cadherin based adhesions serve to connect component cells together. (A similar concern applies to our HeLa cells). Additionally, calcium ions are multipurpose signaling molecules, calcium depletion is therefore expected to impact a host of processes besides cell-cell adhesion (Clapham 2007).
- We do not know the extent to which cell-substrate adhesion in our cells is impacted by the absence of collagen, nor do we know the entire cell-surface receptor profile of our MDCK cells. In agreement with previous work, we observe that cell spreading is diminished in the absence of collagen and more severely on PDMS substrate, but we do not know by how much.

### Cell Crowding Explains the First of Three Architecture Transitions in Cultured Epithelia

In this study we investigated the mechanics of epithelial architecture development in culture. A Mature architecture is characterized by maximum cell density, a polygonal cell arrangement (with respect to the tissue surface) reflected by maximum cell shape regularity, tall cell-cell borders (lateral surfaces), and flat cell apices. Immature architecture is characterized by the minimum density for confluence, irregular cell packing, a lack of cell-cell borders, and the “fried-egg” appearance of cells in which shape (in the apical-basal axis) is determined only by the nucleus. We find that the transitionary morphologies that connect Immature to Mature can be divided into two phases. The first of these, called Intermediate A, arises as a simple consequence of cell crowding. While cell-cell adhesion facilitates the onset of Intermediate A architecture, it may not be a strict requirement; even a non-epithelial cell type (HeLas) can exhibit this architecture at high enough density. The second transitionary phase, Intermediate B, relies on cell-cell adhesion and we therefore propose that it represents the initial apical-basal architecture that is special to epithelia.

The distinction between IntA and IntB architecture corresponds to transitions observed with respect to both the apical-basal axis and the surface of the tissue. With respect to the former, the onset of IntB corresponds to the initial flattening of the apical surface, as revealed by the plot of actin intensity projected across the depth of the tissue. With respect to the latter, the onset of IntB corresponds to a plateau in cell shape regularity. Since cell shape regularity is an indicator of rheological fluidity, this plateau reveals a change in tissue dynamics, from rheologically fluid (characterized by cell rearrangements) to rigid. In our previous work we used live imaging to show that an Intermediate layer undergoes few rearrangements (Dawney et al. 2023). Reevaluation of that layer shows that it is at a density (5.97 x 10^3^ cells/cm^2^) and a circularity (0.84) at which we would expect rigid behavior.

We speculate that the transitions associated with IntB - apical flattening and fluid to rigid tissue dynamics – are functionally linked. Vertex-modeling predicts that the jamming transition occurs at a hallmark shape determined by single-cell mechanical properties, most importantly cell-cell adhesion and contractility (Bi et al. 2015). In epithelia, these two forces are governed largely by an actomyosin belt surrounding each component cell. This belt connects with adherens junctions - specialized epithelial junctions that include E-cadherin - near to the apical surface (Martin and Goldstein 2014). Adhesion is also provided by another type of epithelial junction, the tight junction, which is located apical to the adherens junction (McNeil, Capaldo, and Macara 2006). We showed previously that whereas E-cadherin can be detected in all apical-basal layer architectures, ZO-1, a critical component of the tight junction, is partially observed in Intermediate layers and strongly observed in Mature layers (Dawney et al. 2023). Disruption of ZO-1 in epithelia can lead to increased contractility along apical cell-cell borders (Choi et al. 2016).

In its current state, our biophysical computational model does not include the apical-basal asymmetry in contractility or adhesion conferred by adherens junctions or tight junctions. A goal for future work will be to implement these asymmetries and investigate their contribution to the IntA-IntB and IntB-Mature transitions.

While the regulation of cell polarity - and the impact of polarity on cell shape and tissue organization - have been well-studied in the apical-basal axis, the contribution of cell mechanics in this axis has received less attention. Our novel multimode model for apical-basal shape facilitates the study of mechanical parameters in the establishment and maintenance layered topology. We anticipate that this approach can be used to make additional predictions about the physical parameters that govern epithelial architecture.

## Acknowledgments

We are grateful to the labs of Mark Peifer, Scott Williams, and Holly Lovegrove; the University of Rochester Invertebrate Group; and members of the Bergstralh lab for their questions and comments. This work was supported by an NSF CAREER award (PI: Bergstralh) and NIH Grant R01GM125839 (PI: Bergstralh).

## Competing Interests

The authors declare no competing interests.

## Methods

**Table 1:**
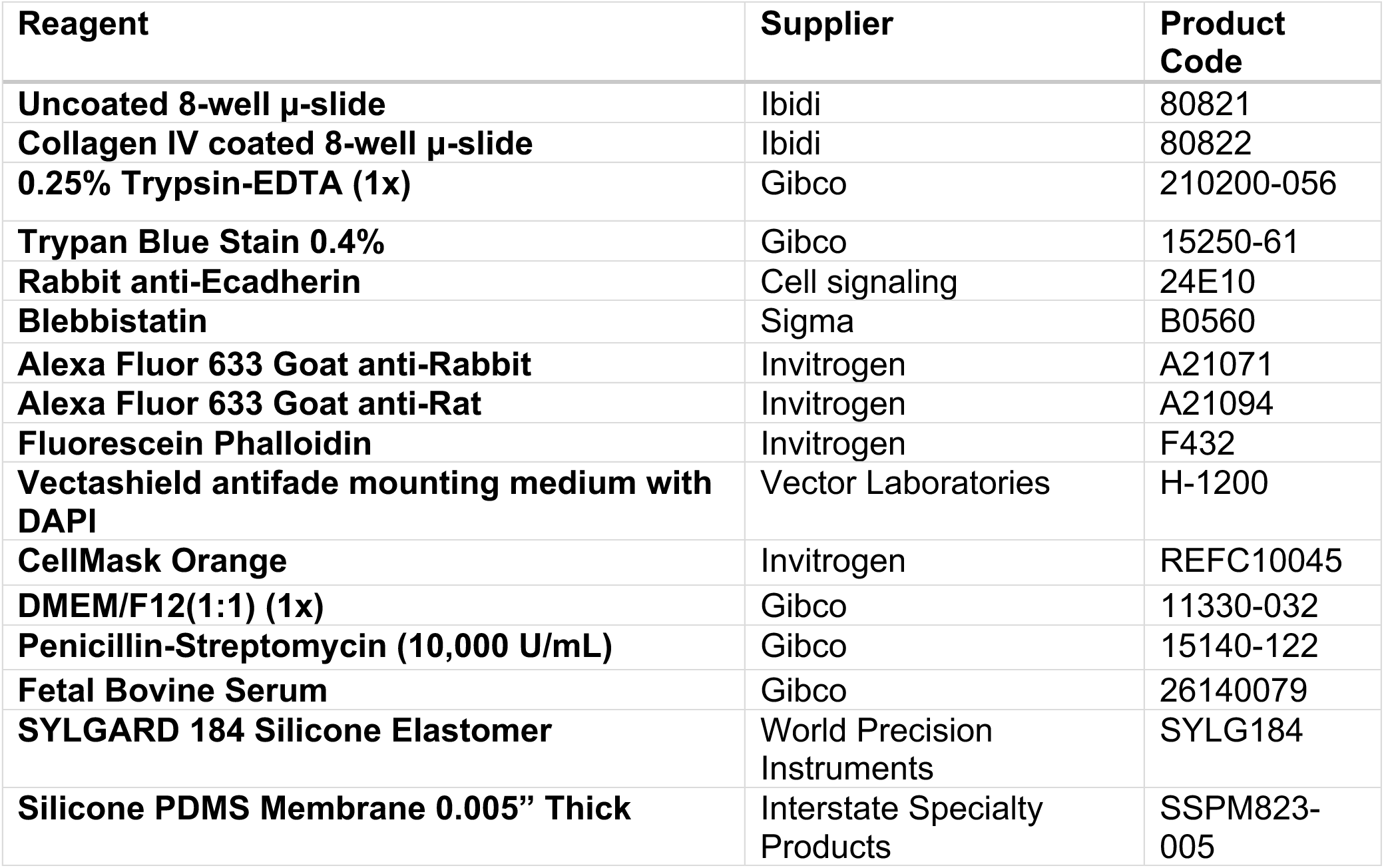
Reagents Used in this Study.

### Computational modeling and Analysis

Computational modeling was undertaken using Python in a Jupyter notebook. Packages used for this model are: Numpy (https://numpy.org/), Matplotlib (https://matplotlib.org/), and NetworkX (https://networkx.org/). Automated Layer Analysis (ALAn) was used to characterize cultured layer architecture. Code is available on the Bergstralh-Lab GitHub repository: https://github.com/Bergstralh-Lab.

The computational model is written dimensionless and is scaled to physical values by interpretation of literature and by comparison to our cultured cell data. The variable parameters values used in model are described in Supplemental Table 2. We calibrated the time scale for our model using the time it takes for two cells to spread: 750 simulated time steps correspond to roughly 1 hour in culture.

**Table 2:** Model variable parameter values used in this Study.

| Variable | Dimensionless Value | Scaled Value | Source |
| --- | --- | --- | --- |
| Area | 1 | $\sim 100 \mu\text{m}^2$ | Observation |
| 2D Area Stiffness | 1 | $\sim 2\text{e}7 \text{ N/m}^2$ | Calculated, Vargas et al 2020 Biophys J (by analogy) |
| Cortical Stiffness | 0.0005 | $\sim 0.1 \text{ N/m}$ | Bruckner et al 2015 Biochim Biophys Acta |
| Cell Cell Adhesion Strength | 0.001 | $\sim 0.2 \text{ N/m}$ | Yeh-Shiu Chu et al 2004 JCB |
| Cell Substrate Adhesion Strength | 0.1 | $\sim 20 \text{ N/m}$ | Gallant, Michael and Garcia 2005 |
| Cell-Cell Interaction Distance | 0.16 | $\sim 1.6 \mu\text{m}$ | Estimate |
| Cell-Substrate Interaction Distance | 0.1 | $\sim 1 \mu\text{m}$ | Estimate |
| Spreading Strength | 0.008 | $\sim 16 \mu\text{N}$ | Observation |

To set a nonlinear scaling between cell-substrate adhesion and cell spreading force, we used a power law. In our nonlinear scaling, spreading force fraction is proportional to the adhesive force fraction to the one fifth.

Division was implemented by having cells double in size over the final hour of their cell cycle before splitting into two daughter cells. The division axis is determined by the top and bottom (in z) cell nodes and produce two daughter cells with a starting cell cycle value taken from a random uniform distribution across the first four hours.

### Cell culture

Cells cultured using standard methods in DMEM supplemented with 10% FBS and 1% Pen/Strep. For standard monolayer culture experiments, cells were passaged one day prior to seeding on 8 well Ibidi chamber slides (uncoated or collagen coated). Media was changed every 24 hrs.

For culture on PDMS membranes, we first cut out a strip of 0.005” thick PDMS membrane the size of a coverslip. Membranes were dipped in methanol to reduce static and placed on a coverslip to allow to dry. A 1 cm by 1 cm culture area was made by curing Sylgard 184 in a petri dish, and cutting a thick walled barrier to mimic the Ibidi culture wells. Slides were kept in a sterile plastic tub and allowed to culture for the appropriate time frame.

To grow MDCK cell colonies, cells were plated at an extremely low density of ∼50 cells onto a 1 cm^2^ collagen coated Ibidi culture slide to get isolated seed cells. Cell culture media was changed every two days, and cells were left for 6 days. Colonies like the one shown in Figure 5 were present in roughly 25% of culture wells, while colonies growing at the edge of the growth area were most common.

### Fixation and Immunostaining

Cells were washed with dPBS and fixed using ∼4% formaldehyde, 2% PBS-tween for 10 minutes. Three washes (10 minutes each) in PBS-0.2% Tween were carried out between fixation and staining.

Primary and secondary antibodies were added at 1:500 dilution. FITC phalloidin was used to stain actin. Vectashield plus DAPI was added directly to the wells. For live imaging, cell mask was added at a concentration of 1:1000 just before imaging. HeLa cells were cultured the same as MDCK cells, but were left for up to a week to promote a homeostatic density.

### Calcium Depletion and Blebbistatin Treatment

Cells were plated and grown as previously described to produce primarily Intermediate architectures (Dawney et al, 2023). After 24 hrs, media was removed and cells were washed with PBS, after which cells were incubated for the specified time in magnesium-calcium-free PBS supplemented with 2 mM Magnesium dichloride. We found that for treatments of up to 3 hours in calcium free media, cell monolayers remained intact and mostly confluent, though there were occasionally cell scale gaps that would develop in the monolayer. These small gaps are unlikely to affect classification by ALAn and are unlikely to impact average architecture of the tissue.

For blebbistatin treated monolayers, cells were grown to an Intermediate architecture and treated with 50 µg/ml of blebbistatin or the equivalent amount of DMSO as a control for the specified time. After treatments cells were fixed and stained as per the above protocol.

### Imaging

For live cell imaging, we used a Leica SP5 Confocal microscope using a 40x/1.25 HCX PL APO oil objective. Confocal stacks were taken at 10-minute intervals.

Fixed tissue was imaged using an Andor Dragonfly Spinning Disk Confocal microscope with a 40x/1.15 water objective. Confocal stacks were taken beginning beneath the bottom of the chamber slide where no actin signal could be detected and ending above the layer, when no more actin signal could be detected, taken at a z-spacing of 0.23 µm. Images were segmented in Imaris, and the corresponding image and .csv files containing nuclear positional information were used with ALAn as previously described. ALAn was used for all architecture classification. Cells cultured on PDMS membranes did not grow to confluence, so regions of 100 pixels x 100 pixels (∼60 µm x 60 µm) were analyzed using ALAn instead of the entire field of view.

To image cell colonies we used a Leica M165 FC Stereomicroscope with a 1.0X objective and a K3M camera.

## Notes

### Competing Interest Statement

The authors have declared no competing interest.

