## Supplemental Data for "The Mechanical Influence of Densification on Initial Epithelial Architecture"

#### Supplemental Figure Legends

***Supplemental Figure 1: MDCK cells take on distinct organized architectures.*** Confocal micrographs of the epithelial architectures traced to provide outlines in Figure 1.

***Supplemental Figure 2: Active spreading is required in the model to recapitulate live cell behavior.*** A) A representative timepoint from live imaging of MDCK cells upon initial culture plating shows that single cells dropped onto a culture substrate spread. Scale bar = 20 μm. B and C) Images of the stable shapes formed by single simulated cells without (B) and with (C) active spreading implemented.

***Supplemental Figure 3: The computational model does not predict the emergence of a Mature architecture.*** The ratio of the apical surface length to the basal surface length of MDCK cells (left) grows as a function of density. Dashed line indicates the density transition between Intermediate and Mature architectures. The same ratio for modeled cells (right). Dashed line indicates predicted transition between Intermediate and Mature architectures.

***Supplemental Figure 4: MDCK Colonies densify at the center.*** The distribution of nuclei (DNA staining) in a colony of MDCK cells. This image, also in Figure 4B, is shown here in a heatmap lookup table to emphasize the gradient of cell density starting from the colony center.

***Supplemental Figure 5: Substrate Adhesion.*** A) Cells grown on collagen spread to cover area of the substrate that is ~5 times larger cells than cells grown on a uncoated substrate. Scale bar = 50 μm. B) Cell-substrate and C) cell-cell contact lengths decrease and increase respectively as layers transition from immature and intermediate architectures. D) Cell-substrate connections decrease and D') Cell-cell connections increase as a function of density at all substrate adhesion strengths with a constant spreading force. The dashed line (D') represents the connection percentage required for

Intermediate architectures to arise in the model. E) Model representative displaying the final result from decreasing cell-substrate adhesion. F) Cell-substrate connections decrease and F’) Cell-cell connections increase as a function of density at all substrate adhesion strengths when cell spreading force is scaled linearly with substrate adhesion. The dashed line (F’) represents the connection percentage required for Intermediate architectures to arise in the model. G) Model representative displaying the final result from decreasing cell-substrate adhesion. H) Cell shape regularity (with respect to the tissue surface) is not impacted by the presence of collagen on the substrate. Only Intermediate layers are shown. $p = 0.4692$, Significance was determined using an unpaired two-tailed Student’s t test.

***Supplemental Figure 6: Blebbistatin treatment does not affect Intermediate layer arhitecture, and HeLA cells form Intermediate architectures.*** A) Blebbistatin does not affect the presence Intermediate architectures at the densities tested. B) MDCK cells express E-cadherin at cell-cell borders, while E-cadherin is not expressed in HeLa cells. C) Cell shape regularity (with respect to the tissue surface) is not impacted by Blebbistatin treatment. Only Intermediate architectures are shown. p values left to right: $p = 0.1495$, $p = 0.7866$, $p = 0.8101$. Significance was determined using an unpaired two-tailed Student’s t test.

***Supplemental Figure 7: HeLa cells do not pack regularly in the plane of the epithelium.*** A) HeLa cells exhibit spindle-like cell morphologies even in layers classed as Intermediate by our image analysis pipeline ALAn. Representative image. Scale bar = 20 μm. B) Cell shape regularity (with respect to the tissue surface) is significantly lower in Intermediate A architectures in comparison to Intermediate B architectures. $p < 0.0001$. Significance was determined using an unpaired two-tailed Student’s t test.

**SUPPLEMENTAL FIGURE 1 Cammarota *et al.***

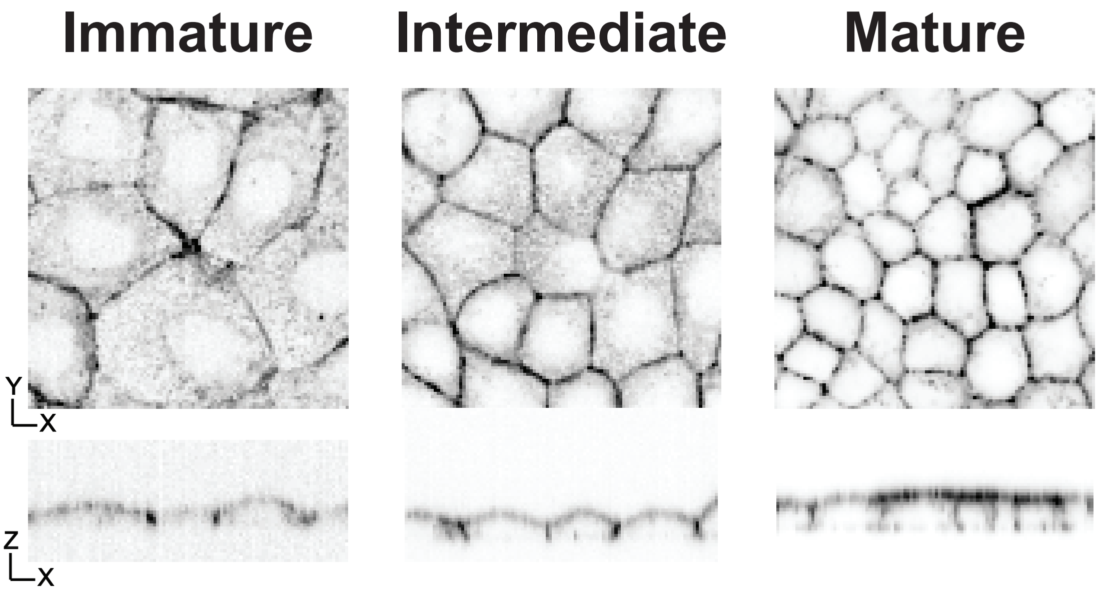

**SUPPLEMENTAL FIGURE 2 Cammarota *et al.***

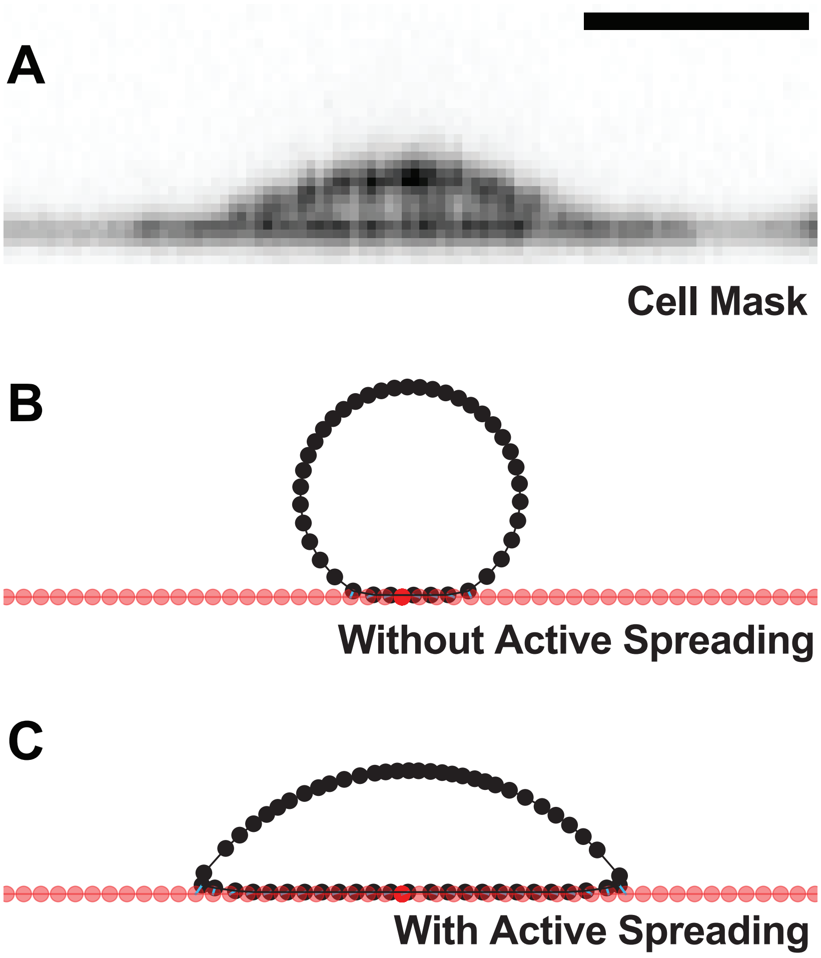

### SUPPLEMENTAL FIGURE 3 Cammarota *et al.*

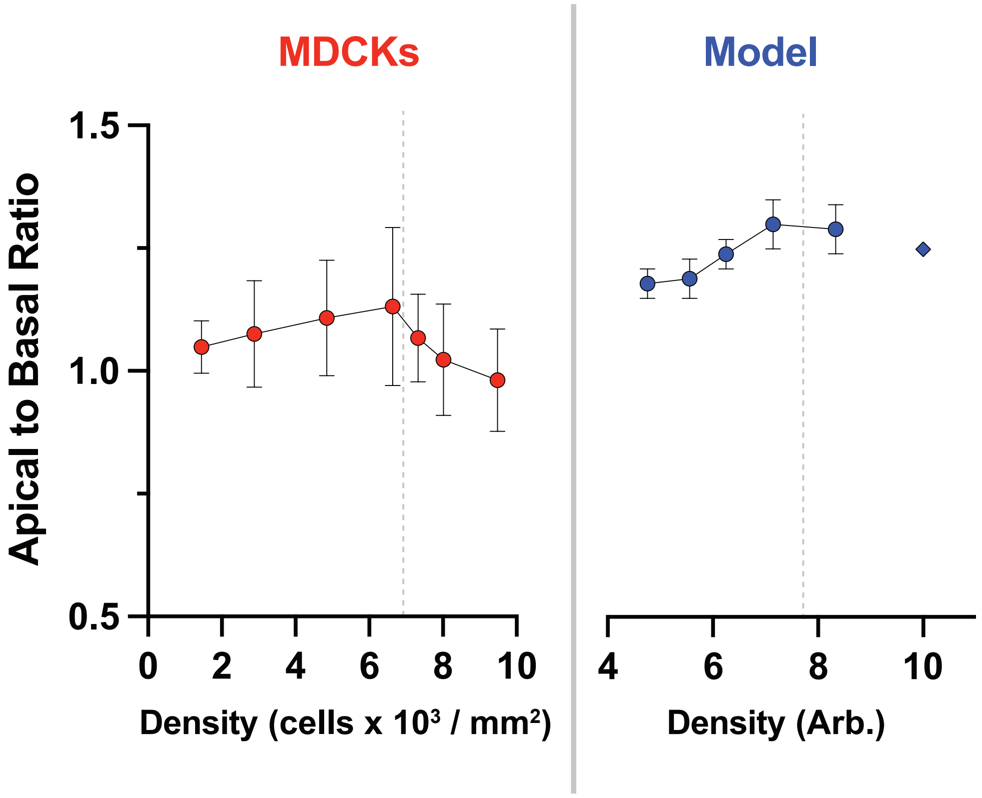

### SUPPLEMENTAL FIGURE 4 Cammarota *et al.*

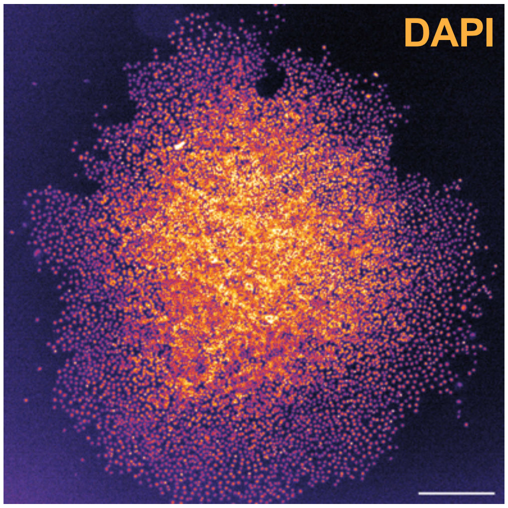

### SUPPLEMENTAL FIGURE 5 Cammarota *et al.*

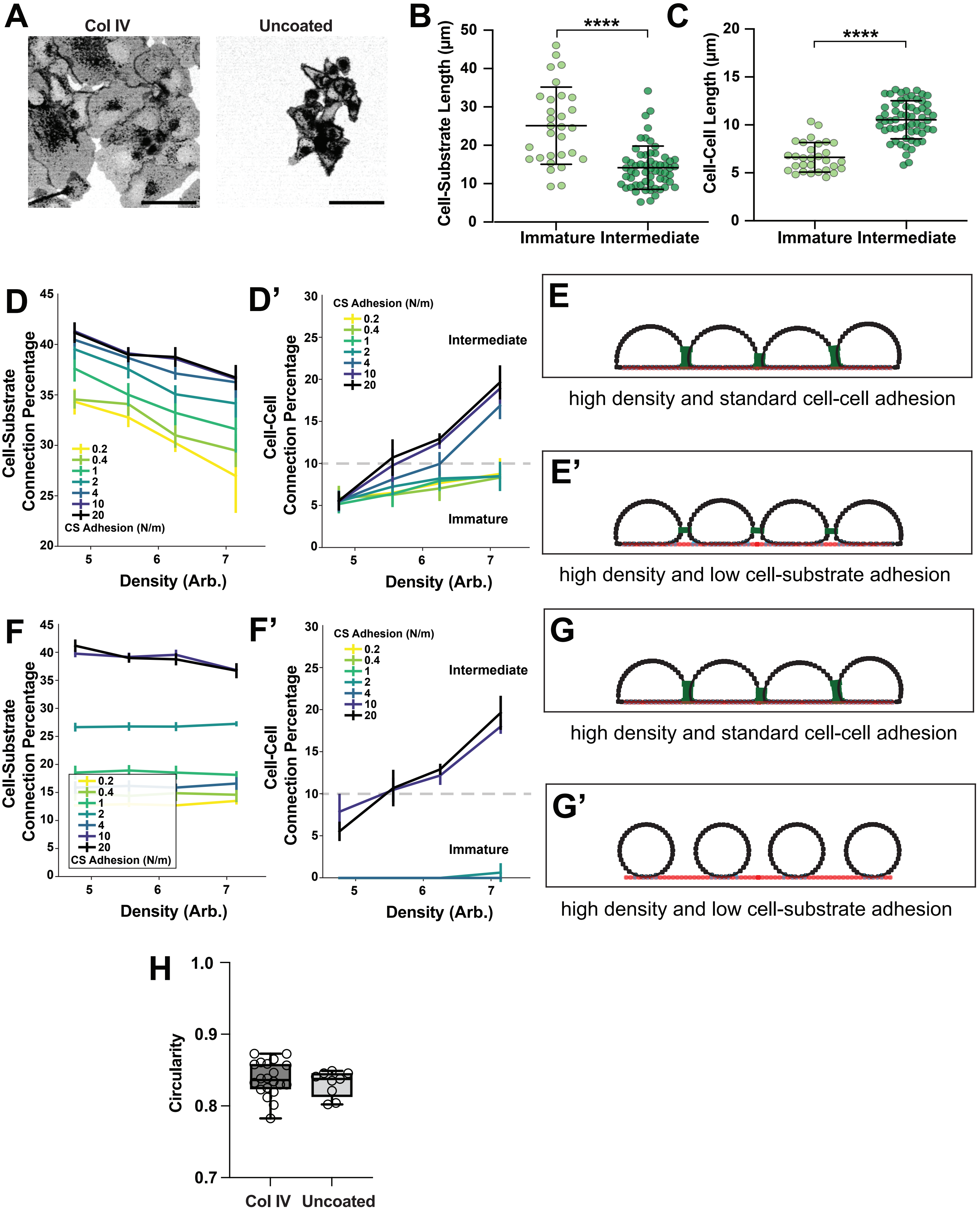

### SUPPLEMENTAL FIGURE 6 Cammarota *et al.*

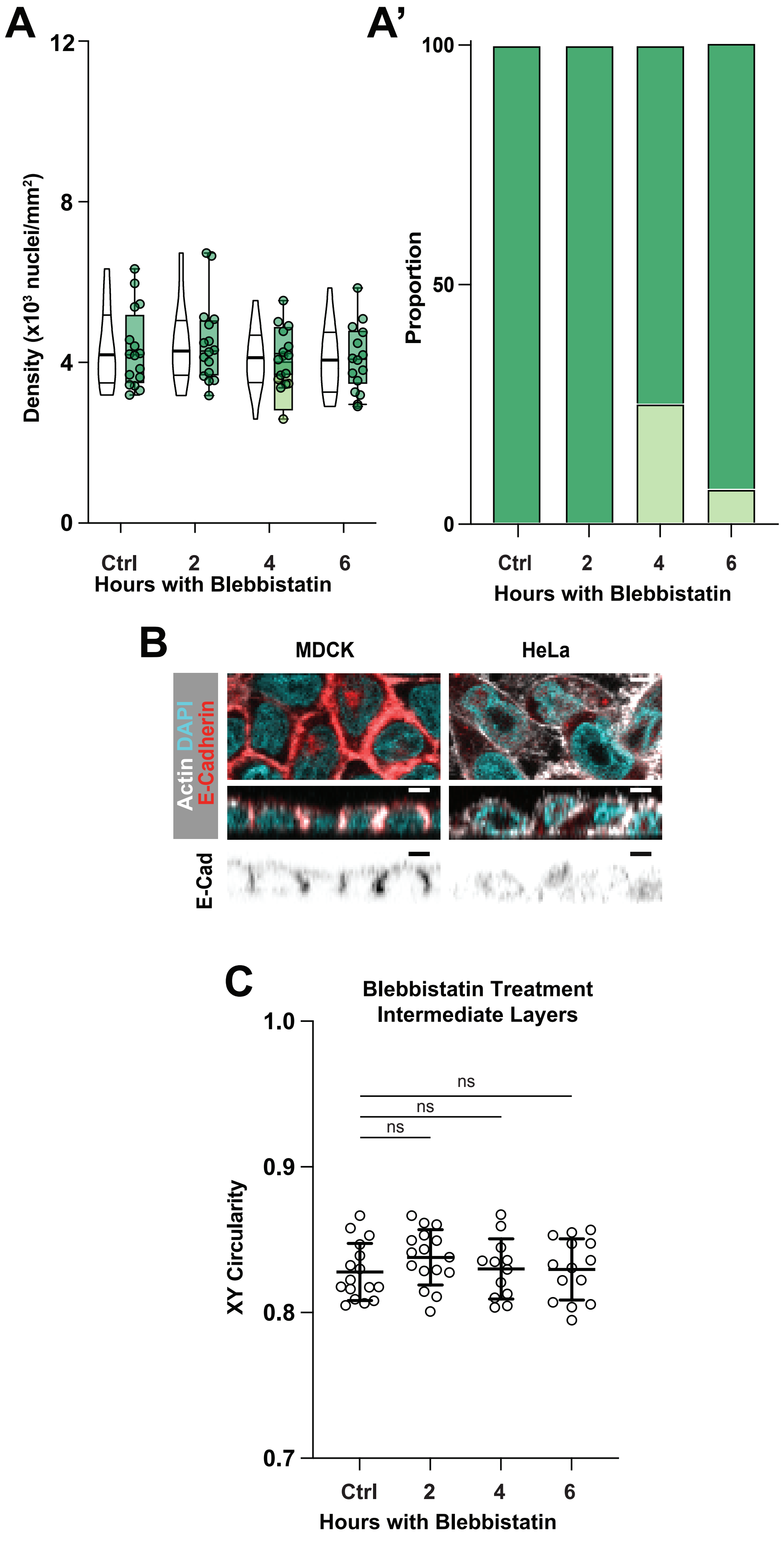

### SUPPLEMENTAL FIGURE 7 Cammarota *et al.*

**A**

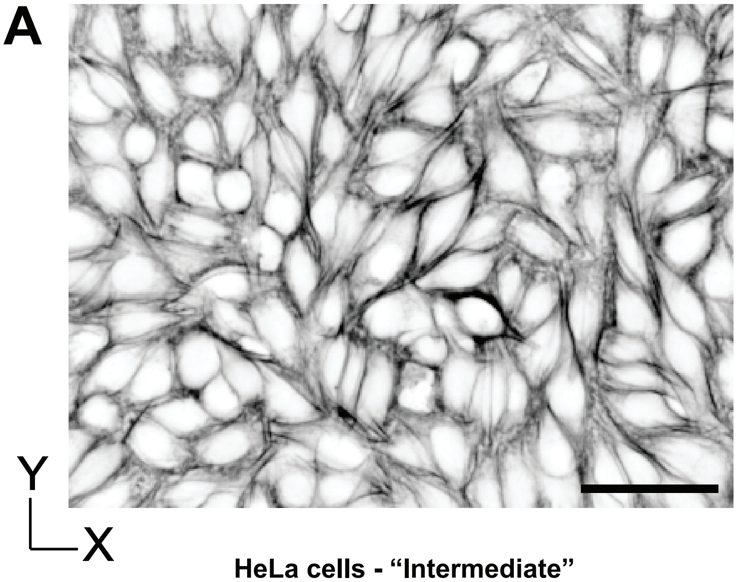


**HeLa cells - “Intermediate”**

**B**

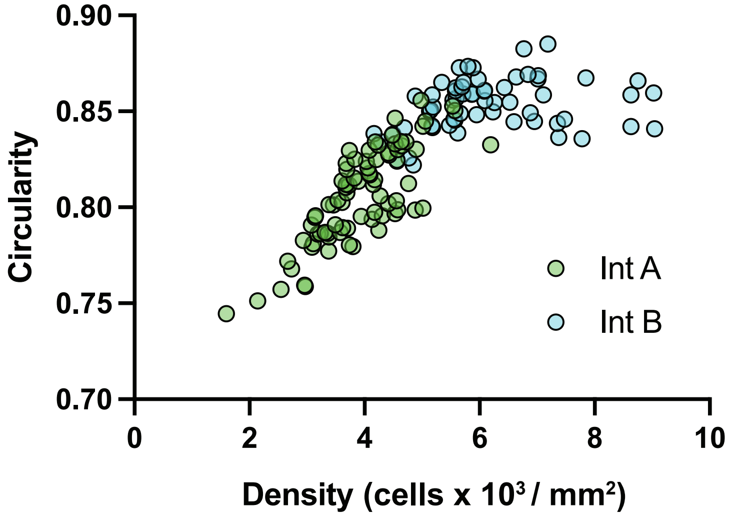
